# Identification of cerebellar afferent projections in a nonhuman primate using CAV-2 vectors

**DOI:** 10.1101/728709

**Authors:** Iria Gonzalez Dopeso-Reyes, Felix Junyent, Nadine Mestre-Francés, Amani Whebi, Bertrand Beucher, Eric J Kremer

## Abstract

Until relatively recently, the cerebellum was considered primarily as modulator of fine motor functions. This has changed during the last 20 years, and now the cerebellum has been shown to be involved in learning and cognition. With this renewed interest comes the need to better understand and potentially modify its input-output connections. In this pilot study, we tested the efficacy of a canine adenovirus type 2 (CAV-2) vector in the cerebellum of a nonhuman primate (NHP). Consistent with other reports, we found a preferentially transduction of cells with neuron-like morphology at the site of injection and efficient retrograde transport into several structures in the midbrain. These data will help identify cerebellar circuits and may lay the foundation for studies of human pathologies, such as ataxias, autism, and schizophrenia.

## Introduction

For most of the 20th century the cerebellum was considered primarily as a modulator of motor functions, and therefore most studies focused on its role in balance, posture and motor control. However, this idea has gradually changed (reviewed by De Smet et al., 2013, Schmahmann, 2018), and now it is thought that the cerebellum plays an important role in language, and in cognitive and affective functions (reviewed by Buckner, 2013). Although it is thought that the cerebellum has maintained most of its neuronal structure and microcircuits throughout vertebrate evolution, its relative size and gross anatomy has changed considerably (Voogd and Glickstein, 1998, reviewed by Llinas et al., 2004, Koibuchi, 2013). This variation in size suggest a relation between size and functional specialization. In humans the cerebellum constitutes about 10% of the total brain weight, but due to the densely packed small granule cells of the cerebellar cortex it contains ∼80% of the total brain neurons (reviewed by Koibuchi, 2013, Essen et al.,2018).

Tract-tracing methods using traditional neuronal tracers have been used to investigate the connections between the cerebellum and the rest of the brain. These methods helped identify anterograde, retrograde or bidirectionally circuits depending on the uptake mechanism or molecular weight (reviewed by Lanciego and Wouterlood 2011). During the last decade viral vectors have been used successfully to unravel and selectively modify brain connections. Viral vectors are powerful tools to study the function of a gene of interest, neurocircuits, structure-function, disease modeling, and therapeutic gene transfer (Wouterlood et al., 2014; Nassi et al., 2015, Mestre-Frances et al., 2018, del Rio et al., 2019, Lasbleiz et al., 2019). Viral vectors expressing reporter proteins can be combined with other technologies such as optogenetics or chemogenetic to better understand brain function (Adamantidis et al., 2015; Urban and Roth, 2015).

Among the vectors used in the CNS, those based on canine adenovirus type 2 (CAV-2), are powerful option to study neuronal circuitry and function (reviewed by del Rio et al. 2019). Studies in the rodent, as well as the dog brain demonstrated that CAV-2 vectors preferentially transduce neurons, lead to widespread distribution via axonal transport (retrograde transport), and are capable of maintaining long-term transgene expression (Soudais et al., 2001, Soudais et al., 2004, Hnasko et al., 2005, Cubizolle et al., 2014, Beier et al., 2015, Schwartz et al., 2015, Salinas et al., 2017, Hirschberg et al., 2017). In the NHP brain, CAV-2 vectors have also been remarkably efficient in the context of preferential transduction of neurons and retrograde transport (Mestre-Frances et al., 2018, Lasbleiz et al., 2019).

Due to the dearth of information concerning cerebellar circuitry in the NHP, and its possible roles in cognitive and affective functions, we assayed the tropism and efficacy of helper-dependent (HD) CAV-2 vectors in the Microcebus murinus (M. murinus). M. murinus is a NHP from Madagascar that is readily bred in captivity, can live >10 years, and naturally shows occasional age-related signs of neurodegenerative disease (Verdier et al.,2015). We report robust CAV-2 vector efficacy at the site of injection as well as in nuclei that project to the injected area.

## Material and methods

One adult male M. murinus was used in this study. Animal handling was conducted in accordance with the European Council directive (2010/63/EU) as well as in agreement with the Society for Neuroscience Policy on the Use of Animals in Neuroscience Research. The experimental design was approved by the Ethical Committee for Animal Testing of the University of Montpellier. The M. murinus colony is housed at the primate facility at the University of Montpellier (license approval n°34-05-026-FS.

### Vector

Construction, purification, and storage of the helper-dependent (HD) CAV-2 vector expressing GFP (HD-GFP) has been described (Soudais et al., 2004). Briefly, HD-GFP contains a cytomegalovirus (CMV) early promoter driving expression of the (enhanced) green fluorescent protein (GFP), followed by a simian virus 40 (SV40) polyA signal. The ∼30 kb HD vector genome does not contain CAV-2 coding regions.

### Stereotaxic Surgery

For surgery, the M. murinus was anesthetized with 80 mg/kg ketamine and10 mg/kg xylazine by intramuscular injections resulting in deep anesthesia. The animal was positioned in the stereotaxic frame; the coordinates for the cerebellum were taken from the M. murinus brain atlas (Bons et al., 1998), - 6.5 mm anteroposterior (AP), - 6 mm dorsoventral (DV) and ± 1 mm mediolateral (ML) from bregma. The M. murinus received two deposits of 1 x109 physical particles of HD-GFP/coordinate through the same needle track (1 μl/coordinate) using a 10 µl Hamilton syringe. HD-GFP was injected at a rate of 0.5 μl/min. Once the deposit was completed, the needle was left in place for 5 min. before withdrawal to minimize vector leakage through the injection tract. The skin was sterilized, sutured and the animal was monitored until recovery from anesthesia. After surgery, the M. murinus was isolated in cage with food and water ad libitum for 24 h.

### Tissue processing and image acquisition

Two weeks post-surgery the M. murinus was anesthetized with ketamine (80 mg/kg) and perfused transcardially with heparinized saline buffer (0.9%) followed by a fixative solution containing 4% paraformaldehyde (PFA) and phosphate buffer saline (PBS), pH 7.4. The brain was post-fixed in 4% PFA and soaked in 30% sucrose/PBS at 4°C for at least 24 h. The brain was embedded in OCT and cut rostro-caudally (50-μm-thick sections) and mounted with Vectashield (Vector Laboratories, H-1000). HD-GFP-mediated GFP expression was detected using a Hamamatsu Nanozoomer with a 40× objective. Background autofluorescence was subtracted using ImageJ. Figures set up was done using Adobe Illustrator CS6.

## Results

To investigate the efficacy of CAV-2 vector tropism and efficacy in the cerebellum and in neurons projecting to the cerebellar cortex, we injected HD-GFP, a helper-dependent CAV-2 vector expressing GFP. Two weeks post-injection, the M. murinus was perfused and the brain was prepared to study the biodistribution GFP expression. Fifty-micron-thick sections were screened by epifluorescent microscopy to identify the regions containing cells expressing GFP (Figure 1). At the site of injection, the granular and the molecular layer (Figure 1B-D) contained numerous GFP+ fibers and somas (Figure 1C). GFP was not observed above the injection site along the needle tract (Figure 1A). Adjacent regions of the cerebellar cortex did not contain GFP+ cells in the molecular or granular layer. At the same level, we observed intense GFP expression in the deep cerebellar nuclei in both hemispheres (Figure E-G). We found numerous GFP+ fibers and some cells in the medial areas of the nucleus interpositus anterior cerebelli (Figure 1F) and in the nucleus mediallis cerebelli (Figure 1G). GFP expression was absent in the nucleus lateralis cerebelli (Figure 1E).

**Figure 1:**
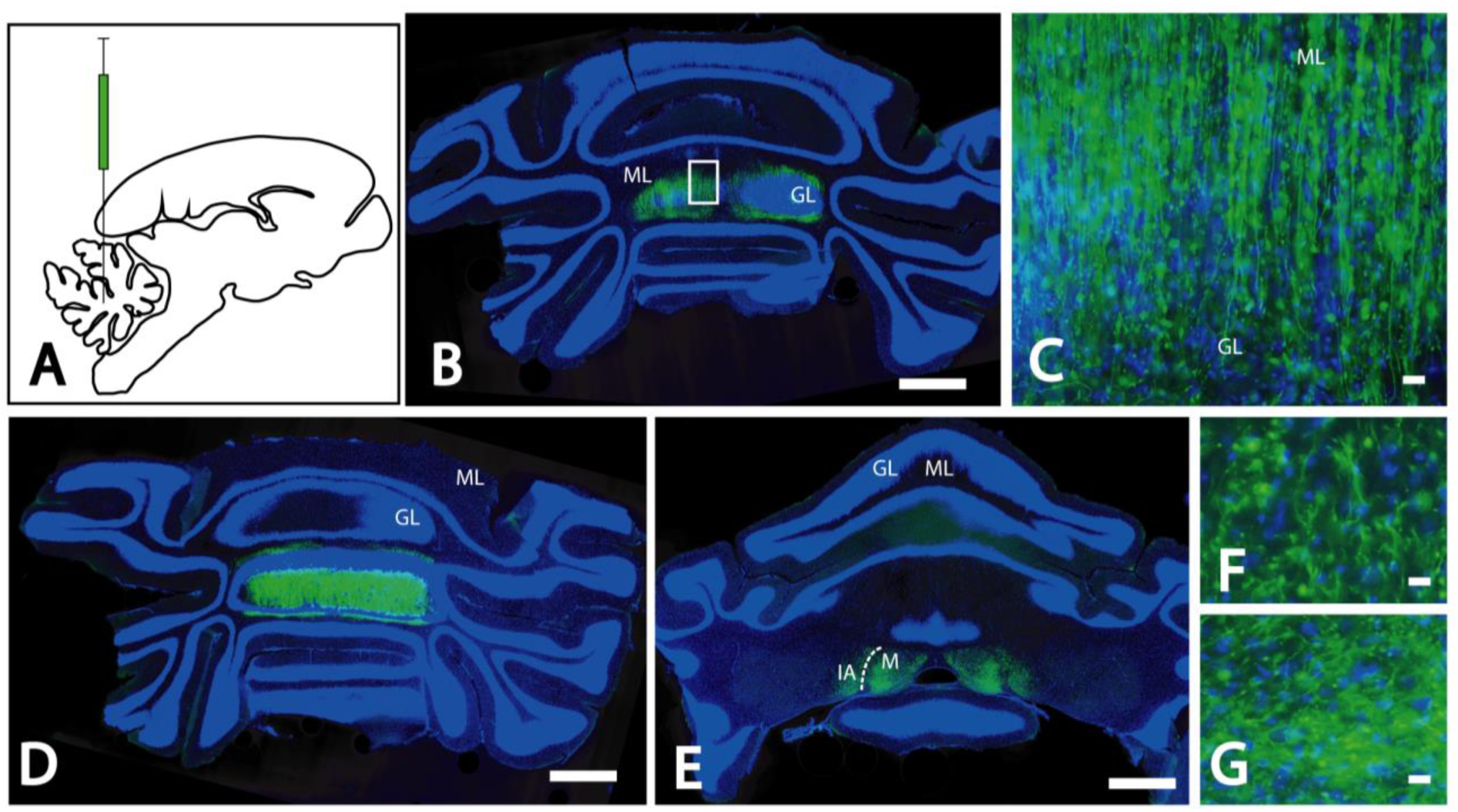
Injection site and GFP expression through the site of the CAV-2 injection. A. Sagittal scheme of the M. murinus showing the site of injection in the cerebellum. B. Coronal section of the cerebellum showing the injection site in both hemispheres of the cerebellum. Nuclei are stain in blue with DAPI showing the histology of the cerebellum, both layers GL and ML show different intensity of GFP expression. C. Detail of B, showing the presence of GFP in numerous fibers in the GL and ML and also numerous somas in the granular layer. D. Coronal section showing the injection site at a different level. Intense expression of GFP in both GL and ML. E. Coronal section showing the injection site at the level of the deep cerebellar nuclei. Presence of GFP expression in IA and M nuclei. F. Detail of the GFP expression in the fibers of the IA. G. Detail of the GFP expression in the fibers of the M. Calibration bars: B, C-D 1 mm, C, F and G 10 µm.

We also found GFP+ cells located in distal cortical areas of the cerebellum (Figure 2A) with morphology consistent with Purkinje cells. Some of these had a weak GFP signal in the soma, but not in the dendritic tree located in the molecular layer. GFP+ cells were located in several nuclei of the medulla oblongata (Figure 2B). In the dorsal areas of the medulla, neurons were GFP+ in the nuclei vestibularis lateralis and medialis, as well as in the nucleus tractus spinales nervi trigemini (Figure 2B). Both nuclei showed numerous GFP+ somas (Figure 2B-C). We also found GFP+ cells in the reticular formation, in the nucleus reticularis magnocellular and in the paramedianus (Figure 2B and 2D). There were several GFP+ cells in the reticular formation (Figure 2D). The inferior olive also showed some intense GFP+ cells, those mainly in its caudal region and scattered GFP+ fibers (Figure 2E). By contrast, we did not observe GFP+ fibers in the main tracts of the medulla, the pyramidal tracts, lemniscus medialis and the fasciculus longitudinalis medialis (Figure 2B).

**Figure 2:**
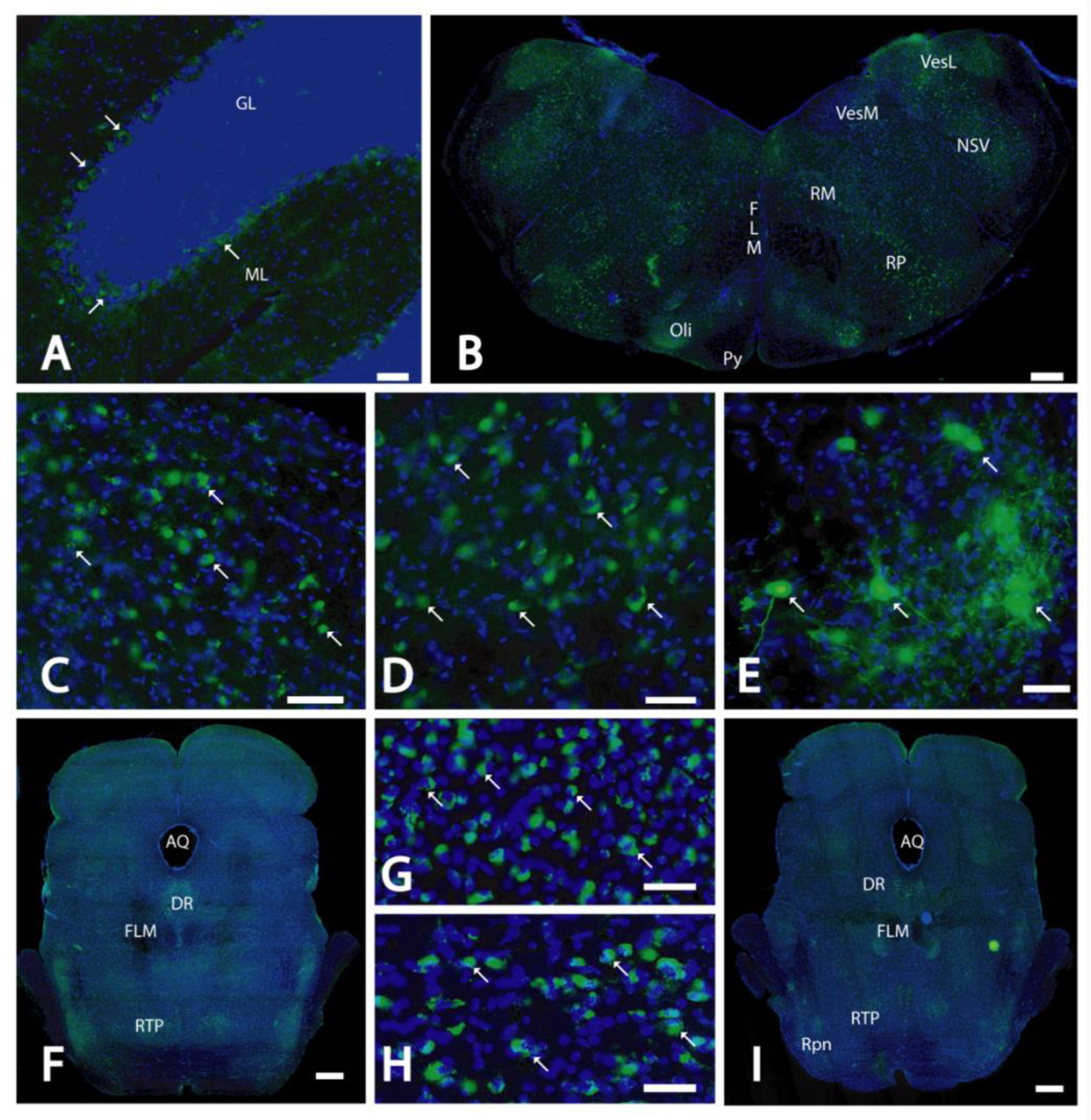
Coronal sections showing the GFP expression showing the GFP expression. A. Detail showing the presence of GFP in the soma of Purkinje cells (arrows) in some areas of the cerebellar cortex, notice the lack of GFP in the dendritic tree and in the other layers, GL and ML. B. Coronal section of the medulla oblongata showing the presence of different neuronal populations expressing GFP. C. High magnification of the VesM showing the presence of GFP in some of the cell somas (arrows). D. High magnification demonstrating the presence of GFP in some of the cell somas (arrows). E. High magnification of the Oli where we show some GFP+ fibers and some of the somas (arrows) with an intense expression of GFP. F. Coronal sections of the pons showing the different neuronal population expressing GFP. G. High magnification of GFP+ somas (arrows) in the DR. H. High magnification showing the GFP expression in several somas (arrows) of the RTP. I. Coronal section of more rostral section of the pons showing the different GFP+ neuronal populations. Calibration bars: A 50 µm, B, F and I 1 mm, C-E, G-H 20 µm.

In more rostral regions, GFP+ cells were detected in the pontine reticular formation nuclei (Figures 2F-I), in the nucleus dorsalis raphes (Figures 2F-G and 2I), in the nucleus reticular tegmenti pontis (Figures 2F, 2H-I), and in the nucleus raphes pontis (Figure 2I). The pontine and medullar nuclei also showed scattered GFP+ fibers (data not shown).

## Discussion

Gene therapy for neurodegenerative disorders will be irreversible – in other words, once a viral vector is injected into the brain there will be no way to remove it. Therefore, this therapeutic approach creates the need to address vector tropism, feasibility, efficacy, safety, and duration of expression in an animal that closely resembles humans. In this study, we evaluated the efficacy of CAV-2 vectors in the M. murinus cerebellum and found robust transgene expression and biodistribution. Neurons in distal structures can readily take up CAV-2 vectors if the neuron expresses the coxsackievirus adenovirus receptor (CAR) (Salinas et al., 2017, del Rio et al. 2019). It is possible that more vector is taken up by some structures due to the density of their projections near the injection site and their level of CAR expression in the axons. It is unlikely that the ∼100 nm, neutral-charged CAV-2 capsid (Schoehn et al., 2008) diffuses significantly in the brain parenchyma, and therefore neurons are capable of continually taking up vector particles deposited at the injection site for an extended period, possibly for several days. We previously demonstrated that the transcriptional changes after HD-GFP injection in the M. murinus are likely due to detection of the free ends of the double-stranded DNA vector genome. Nonetheless, long-term transgene expression demonstrates that this transcriptional response does not lead to a notable adaptive immune response.

After the injection in cerebellar cortex, we have found GFP+ neurons and fibers in the molecular and granular layers of the cerebellar cortex, in the deep cerebellar nuclei, several precerebellar nuclei and the inferior olivary complex. These structures and cells project to the cerebellum in rodents and primates (reviewed by Glickstein et al.2011, Llinas et al., 2004, Barmack and Yakhnitsa, 2013; Voogd et al., 2013). The molecular layer showed a high number of GFP+ fibers, possibly corresponding with the climbing fibers from the inferior olive and also parallel fibers from the granular cells of the cerebellar cortex. In accordance with this, we observed GFP+ neurons in the inferior olive, origin of the climbing fibers and GFP+ somas in the granular layer origin of the parallel fibers.

The granular layer also contained numerous fibers GFP+. Considering the location of those fibers and also the GFP+ somas observed in different precerebellar nuclei, we believe that they are mossy fibers originating in the nuclei vestibularis lateralis and medialis, nucleus tractus spinales nervi trigemini and reticular formation in the medulla oblongata and the pons. Two deep cerebellar nuclei, nucleus interpositus anterior cerebelli and the nucleus mediallis cerebelli showed numerous GFP+ fibers consistent with the mossy fibers originated in different precebellar nuclei.

The paucity of animal models that faithfully recapitulate human brain diseases has slowed the development of therapies. Transgenic mice have often poorly mimicked human diseases, possibly due to species-specific differences in disease mechanisms and in the anatomical organization (Herodin et al., 2005, Tuszynski et al., 2002). Studies using NHPs should be readily translatable to humans. However, the number of species of NHPs available for biomedical research, such as baboons, macaques or marmosets, is limited, under increased regulatory regulations, and expensive (Languille et al., 2012). Of note, M. murinus are one of the few NHPs approved by current European regulation, and self-sustaining breeding colonies can provide sufficient number of animals for trials that require large cohorts. M. murinus is small primate, whose brain has a structure and organization comparable to that of the human brain.

In conclusion, our results in the M. murinus cerebellum have notable fundamental and clinical implications in the context of gene transfer in the NHP brain. Numerous diseases may be amendable to therapy and modeling using CAV-2 vectors, regardless of whether they affect specific nuclei or the entire brain.

## Abbreviatures

GL: Granular layer of the cerebellar cortex
ML: Molecular layer of the cerebellar cortex
IA: Nucleus interpositus anterior cerebelli
M: Nucleus medialis cerebelli
VesM: Nucleus vestibularis medialis
VesL: Nucleus vestibularis lateralis
NSV: Nucleus tractus spinalis nervi trigemini
RM: Nucleus reticularis magnocellularis
RP: Nucleus reticularis paramedialis
FLM: Fasciculus longitudinalis medialis
Oli: Nucleus olivaris inferior
Py: Tractus pyramidalis
RL: Nucleus reticularis lateralis
LM: Lemniscus medialis
AQ: Aquaductus
DR: Nucleus dorsalis raphes
RTP: Nucleus reticularis tegmenti pontis
nP: Nuclei ponti
Rpn: Nucleus raphes pontis
NR: Nucleus ruber
4th: V 4th ventricle

## Author Contributions

NMF and EJK contributed to the conception and design of the study; FJ performed image acquisition; AW and IGDR analyzed the images; BB provided the vectors, IGDR and EJK wrote the manuscript with contributions from all co-authors.

## Acknowledgments

We thank EKL members for constructive comments during the course of this study. We thank Réseau d’Histologie Expérimental de Montpellier, Montpellier Ressources Imagerie (ANR-10-INBS-04, “Investment for the future”), and the Réseau des animaleries de Montpellier.

## References

Adamantidis, A., Arber, S., Bains, J. S., Bamberg, E., Bonci, A., Buzsáki, G., et al. (2015). Optogenetics: 10 years after ChR2 in neurons - Views from the community. Nat. Neurosci. 18, 1202–1212. doi: 10.1038/nn.4106

Barmack NH and Yakhnitsa V. (2013). Vestibulocerebellar Connections in Handbook of the Cerebellum and Cerebellar Disorders, Manto M, Gruol DL, Schmahmann JD, Koibuchi N, Rossi F (eds.), pp 357-375. DOI 10.1007/978-94-007-1333-8_62, Springer Science+Business Media Dordrecht

Beier KT, Steinberg EE, DeLoach KE, Xie S, Miyamichi K, Schwarz L, Gao XJ, Kremer EJ, Malenka RC, Luo L. (2015). Circuit Architecture of VTA Dopamine Neurons Revealed by Systematic Input-Output Mapping. Cell. 162(3):622–34. doi: 10.1016/j.cell.2015.07.015.

Buckner RL. (2013). The Cerebellum and Cognitive Function: 25 Years of Insight from Anatomy and Neuroimaging. Neuron. 80(3):807–15. doi: 10.1016/j.neuron.2013.10.044.

Cubizolle, A., Serratrice, N., Skander, N., Colle, M. A., Ibanes, S., Gennetier, A., et al. (2014). Corrective GUSB transfer to the canine mucopolysaccharidosis VII brain. Mol. Ther. 22, 762–773. doi: 10.1038/mt.2013.283

De Smet HJ, Paquier P, Verhoeven J, Mariën P. (2013). The cerebellum: its role in language and related cognitive and affective functions. Brain Lang. 127(3):334–42. doi: 10.1016/j.bandl.2012.11.001.

del Rio D, Beucher B, Lavigne M, Wehbi A, Gonzalez Dopeso-Reyes I, Saggio I and Kremer EJ (2019) CAV-2 Vector Development and Gene Transfer in the Central and Peripheral Nervous Systems. Front. Mol. Neurosci. 12:71. doi: 10.3389/fnmol.2019.00071.

Glickstein M, Sultan F, Voogd J. (2011). Functional localization in the cerebellum. Cortex. 47(1):59–80. doi: 10.1016/j.cortex.2009.09.001

Hérodin F, Thullier P, Garin D, Drouet M. (2005). Nonhuman primates are relevant models for research in hematology, immunology and virology. Eur Cytokine Netw 16(2):104–116

Hirschberg S, Li Y, Randall A, Kremer EJ, Pickering AE. (2017). Functional dichotomy in spinal-vs prefrontal-projecting locus coeruleus modules splits descending noradrenergic analgesia from ascending aversion and anxiety in rats. Elife. 6. pii: e29808. doi: 10.7554/eLife.29808.

Hnasko, T. S., Sotak, B. N., and Palmiter, R. D. (2005). Morphine reward in dopamine-deficient mice. Nature 438: 854–857.

Koibuchi N. (2013). Animal Models: An Overview in Handbook of the Cerebellum and Cerebellar Disorders, Manto M, Gruol DL, Schmahmann JD, Koibuchi N, Rossi F (eds.), pp1427- DOI 10.1007/978-94-007-1333-8_62, Springer Science+Business Media Dordrecht

Lanciego JL, Wouterlood FG. (2011). A half century of experimental neuroanatomical tracing. J Chem Neuroanat. 42(3):157–83. doi: 10.1016/j.jchemneu.2011.07.001.

Languille S, Blanc S, Blin O, Canale CI, Dal-Pan A, Devau G, Dhenain M, Dorieux O, Epelbaum J, Gomez D, Hardy I, Henry PY, Irving EA, Marchal J, Mestre-Francés N, Perret M, Picq JL, Pifferi F, Rahman A, Schenker E, Terrien J, Théry M, Verdier JM, Aujard F. (2012). The grey mouse lemur: a non-human primate model for ageing studies. Ageing Res Rev. 11(1):150–62. doi: 10.1016/j.arr.2011.07.001.

Lasbleiz, C., Mestre-Frances, N., Devau, G., Luquin-Piudo, M., Tenenbaum, L., Kremer, E. J., et al. (2019). Combining gene transfer and nonhuman primates to better understand and treat Parkinson’s disease. Front. Mol. Neurosci. 12:10. doi: 10.3389/fnmol.2019.00010

Llinas RR, Walton KD, Lang EJ. (2004). Cerebellum in The Synaptic Organization of the Brain. Shepherd GM (ed) pp 271–310. DOI: 10.1093/acprof:oso/9780195159561.001.1 Oxford University Press, New York.

Mestre-Francés N, Serratrice N, Gennetier A, Devau G, Cobo S, Trouche SG, Fontès P, Zussy C, De Deurwaerdere P, Salinas S, Mennechet FJ, Dusonchet J, Schneider BL, Saggio I, Kalatzis V, Luquin-Piudo MR, Verdier JM, Kremer EJ. (2018). Exogenous LRRK2G2019S induces parkinsonian-like pathology in a nonhuman primate. JCI Insight. 3(14). pii: 98202. doi: 10.1172/jci.insight.98202

Nassi JJ, Cepko CL, Born RT, Beier KT. (2015). Neuroanatomy goes viral! Front Neuroanat. 2015 Jul 1;9:80. doi: 10.3389/fnana.2015.00080

Salinas S, Junyent F, Coré N, Cremer H, Kremer EJ. (2017). What is CAR doing in the middle of the adult neurogenic road? Neurogenesis (Austin). 4(1):e1304790. doi: 10.1080/23262133.2017.1304790

Schmahmann JD. (2018). The cerebellum and cognition. Neurosci Lett. 688:62–75. doi: 10.1016/j.neulet.2018.07.005.

Schoehn G, El Bakkouri M, Fabry CM, Billet O, Estrozi LF, L. L, Curiel DT, Kajava AV, Ruigrok RW, Kremer EJ. (2008). Three-dimensional structure of canine adenovirus serotype 2 capsid. J Virol. 2008 Apr;82(7):3192–203. doi: 10.1128/JVI.02393-07.

Schwarz LA, Miyamichi K, Gao XJ, Beier KT, Weissbourd B, DeLoach KE, Ren J, Ibanes S, Malenka RC, Kremer EJ, Luo L. 2015. Viral-genetic tracing of the input-output organization of a central noradrenaline circuit. Nature. 524 (7563):88–92. doi: 10.1038/nature14600.

Soudais C, Laplace-Builhe C, Kissa K, Kremer EJ. (2001). Preferential transduction of neurons by canine adenovirus vectors and their efficient retrograde transport in vivo. FASEB J. 15(12):2283–5.

Soudais, C., Skander, N., and Kremer, E. (2004). Long-term in vivo transduction of neurons throughout the rat CNS using novel helper-dependent CAV-2 vectors. FASEB J. 18, 391–393. doi: 10.1096/fj.03-0438fje

Tuszynski MH, Grill R, Jones LL, McKay HM, Blesch A. (2002). Spontaneous and augmented growth of axons in the primate spinal cord: effects of local injury and nerve growth factor-secreting cell grafts. J Comp Neurol. 449(1):88–101.

Urban DJ, Roth BL. (2015). DREADDs (designer receptors exclusively activated by designer drugs): chemogenetic tools with therapeutic utility. Annu. Rev. Pharmacol. Toxicol. 55 399–417. 10.1146/annurev-pharmtox-010814-12480

Van Essen DC, Donahue CJ, Glasser MF. (2018). Development and Evolution of Cerebral and Cerebellar Cortex. Brain Behav Evol. 91(3):158–169. doi: 10.1159/000489943

Verdier JM, Acquatella I, Lautier C, Devau G, Trouche S, Lasbleiz C, Mestre-Francés N. (2015). Lessons from the analysis of nonhuman primates for understanding human aging and neurodegenerative diseases Front. Neurosci. https://doi.org/10.3389/fnins.2015.00064

Voogd J, Glickstein M. (1998). The anatomy of the cerebellum. Trends Neurosci. 21(9):370–5.

Voogd J, Shinoda Y, Ruigrok TJH, Sugihara I. (2013). Cerebellar Nuclei and the Inferior Olivary Nuclei: Organization and Connections in Handbook of the Cerebellum and Cerebellar Disorders, Manto M, Gruol DL, Schmahmann JD, Koibuchi N, Rossi F (eds.). pp 377–436. DOI 10.1007/978-94-007-1333-8_62, Springer Science+Business Media Dordrecht

Wouterlood FG, Bloem B, Mansvelder HD, Luchicchi A, Deisseroth K. (2014). A fourth generation of neuroanatomical tracing techniques: exploiting the offspring of genetic engineering. J Neurosci Methods. 235:331–48. doi:10.1016/j.jneumeth.2014.07.021

